# Developing Interferon-β as a safe in-vivo experimental-medicine model of human inflammation

**DOI:** 10.1101/2025.02.18.638888

**Authors:** Eva Periche-Tomas, Claire Maclver, John Underwood, Helena Leach, Claudia Bone, Simon Jones, Barbara Szomolay, Kathy Triantafilou, Neil A. Harrison

**Affiliations:** Cardiff University Brain Research Imaging Centre (CUBRIC), School of Psychology, UK; Hodge Centre for Translational Neuroscience, School of Medicine, Cardiff University, UK; School of Medicine, Cardiff University, UK; Division of Infection and Immunity, School of Medicine, Cardiff University, UK; Systems Immunity Research Institute, School of Medicine, Cardiff University, UK

## Abstract

**Background:** Inflammation is increasingly implicated in a wide range of neuropsychiatric disorders ranging from depression through age- and infection-related cognitive decline to dementia. Arguably, the most important evidence supporting an aetiological role for inflammation in these conditions in humans has come from studies of patients receiving IFN- α therapeutically or longitudinal follow-up of patients after naturalistic infections (e.g., hepatitis C). These experimental medicine-type approaches have identified a discrete set of brain regions e.g. amygdala-hippocampus-hypothalamus, insula and anterior cingulate as well as dopamine-rich subcortical structures that are particularly sensitive to systemic inflammation. Coupled with renewed interest in developing novel immune-targeted therapies for neuropsychiatric disorders as diverse as depression and neurodegenerative disorders, this has highlighted the urgent need for a safe, reliable in-vivo experimental medicine model of inflammation that can be used across the age range. To date, this need has been partially addressed by access to short-acting forms of Interferon-alpha or human (GMP) grade lipopolysaccharide (LPS). However, unpegylated IFN-α is no longer commercially available and the costs and cardiovascular monitoring requirements of low-dose (i.e. 0.8-1ng/Kg i.v. endotoxin) LPS are prohibitive and particularly challenging to use in older or more vulnerable populations.

**Aim:** Develop a new experimental model of human inflammation that elicits robust sickness responses within a few hours but with minimal cardiovascular effects, thus avoiding the need for continuous cardiac monitoring and ensuring applicability across diverse experimental contexts and participant groups from the young to the elderly.

**Methods:** Using a randomized, placebo-controlled, repeated measures cross-over design, physiological, behavioural, immune and transcriptomic responses were collected from 30 healthy volunteers (15 young [18-34] and 15 old [60-75]) following both IFN-β (EXTAVIA® [100 µg]) and saline (placebo) injections.

**Results:** IFN-β induced a robust systemic immune response, evidenced by significant increases in temperature, heart rate, immune cell activation (lymphocytes, monocytes and neutrophils) and levels of IFN-β, IL-10 and TNF-α cytokines. These physiological changes were accompanied by significant increases in negative mood, tiredness, tension and sickness symptoms aa well as by a decrease in vigour.

**Conclusions:** FN-β is a safe and robust new experimental model of mild acute inflammation, This minimally invasive and effective design can induce transient changes in systemic inflammation in healthy individuals from 18-75. A more refined, ecologically valid model similar to the mild inflammation typically reported in psychiatric disorders like depression or cognitive impairment.

## Introduction

Converging evidence from epidemiological, experimental and preclinical studies has increasingly implicated inflammation in a wide range of neuropsychiatric disorders ranging from depression to age-related cognitive decline and dementia (Khandaker et al., 2021). Recent experience with COVID has also reminded us that robust host immune responses during severe systemic infections can also result in sustained/persistent cognitive impairment even in previously healthy individuals (Iwashyna et al., 2010; Wood et al., 2025). However, of these approaches, arguably, the strongest evidence for an aetiological role for inflammation in human neuropsychiatric disorders has come from experimental medicine studies. This includes acute immune-challenge models typically undertaken in healthy participants and studies of patients receiving regular IFN-α injections over many months for therapeutic purposes (Capuron et al., 2007; Capuron & Miller, 2004; Harrison et al., 2014, 2015, 2016) which have predominantly focussed on mechanisms of mood and motivational change and to a lesser degree cognitive effects, as well as follow-up studies of patients after naturalistic infections that have had a greater focus on long-term cognitive deficits (Iwashyna et al., 2010; Wood et al., 2025).

Together, these approaches have helped to identify the discrete set of brain regions that appear particularly sensitive to systemic inflammation. This includes regions of the medial temporal lobe such as the amygdala and hippocampus, and the hypothalamus that are integral to stress and memory circuits (Harrison, 2017). The insula, dorsal, posterior and sub-genual cingulate and peri-aqueductal grey (PAG) that form the interoceptive and autonomic control networks, salience and default mode networks (Harrison et al., 2009; Kitzbichler et al., 2021) and dopamine-rich subcortical structures that are critical to motor responses, reward-related behaviours and motivational control (Harrison, 2017).

Furthermore, those studies that have simultaneously investigated the acute and chronic effects of IFN-alpha in the same participants have shown that individuals with stress and reward systems that are most sensitive to acute inflammation also go on to develop the most severe depressive symptoms and motivational impairments during chronic IFN-alpha administration (Capuron et al., 2007; Capuron & Castanon, 2017; Davies et al., 2020; Dowell et al., 2016, 2019). In the context of fatigue-related motivational impairment, preliminary transcriptomic analyses suggest that this sensitivity may be related to early activation of m-TOR pathways (Periche-Tomas et al., 2025)

Together, this suggests that acute and chronic immune challenges recruit and disrupt similar brain networks. Furthermore, it suggests that acute experimental immune challenge models can serve as a reasonable proxy for predicting inter-individual differences in susceptibility to the more severe and potentially long-lasting effects of more intense or chronic inflammation that would be unethical to investigate experimentally outside of a clinically therapeutic context. Coupled with renewed interest in developing novel immune-targeted therapies for neuropsychiatric disorders as diverse as depression and neurodegenerative disorders, this has highlighted the urgent need for a safe, reliable in-vivo experimental medicine model of inflammation that can be used across the age range. To date, this need has been partially addressed by access to short-acting forms of Interferon- α or human (GMP) grade lipopolysaccharide (LPS). However, unpegylated IFN-α is no longer commercially available and the costs and cardiovascular monitoring requirements of low-dose (i.e. 0.8-1ng/Kg i.v. endotoxin) LPS are prohibitive and particularly challenging to use in older or more vulnerable populations.

This study presents a novel model of acute experimental inflammation using IFN-β, designed to achieve a balance between efficacy and minimal invasiveness. Unlike existing models, such as endotoxin challenges, this approach is both less invasive and highly effective. IFN-β’s capacity to elicit a controlled yet robust inflammatory response makes it a versatile tool suitable for experimental applications across diverse populations, including both younger and older individuals.

Type I interferons (IFNs), particularly IFN-α and IFN-β, are crucial antiviral cytokines that regulate inflammatory responses by activating the JAK-STAT pathway via the IFNAR1/IFNAR2 receptor, leading to the induction of interferon-stimulated genes (ISGs) and the secretion of cytokines such as IL-6, TNF-α, and IL-1ra (Kasper & Reder, 2014; Kümpfel et al., 2000). IFN-α was widely used to treat Hepatitis C (McHutchison et al., 1998), and IFN-β is a cornerstone therapy for multiple sclerosis (MS), helping reduce recurrence rates and delaying disability ( Jacobs et al., 1981, 1982; Jacobs et al., 1996).

IFN-α impedes Hepatitis C virus (HCV) replication by inducing an antiviral state in infected and neighbouring cells, enhancing natural killer (NK) cell activity, and disrupting HCV protein processing (Gale & Katze, 1998; Rehermann, 2013; Samuel, 2001). In contrast, IFN-β’s mechanism in MS involves several immunomodulatory effects: inhibiting T-cell activation and proliferation, suppressing pro-inflammatory cytokine production, enhancing anti-inflammatory cytokine production, reducing the expression of adhesion molecules, and promoting regulatory T-cell expansion (Cheng et al., 2015; Kieseier, 2011; Mirandola et al., 2009; Teige et al., 2006; Windhagen et al., 1995).

Peripheral administration of Type I interferons, including IFN-β, can access the brain through various mechanisms: increased BBB permeability, active transport systems, release of inflammatory mediators from cerebral vasculature, and signalling via peripheral nerve fibres (Banks, 2016; Banks & Erickson, 2010; Erickson & Banks, 2018; Goehler et al., 2000; Pan et al., 1997). This is supported by rodent studies showing upregulation of ISGs in the brain following IFN administration (Wang, 2009; Wang & Campbell, 2005). In humans, IFN-α administration has been shown to stimulate IL-6 and TNF-α production in peripheral blood cells, with elevated levels found in the cerebrospinal fluid of patients undergoing treatment (Capuron et al., 2003; Raison et al., 2009; Taylor & Grossberg, 1998).

IFN injections induce a systemic inflammatory response mimicking viral infection, often resulting in flu-like symptoms (Davis et al., 1989; Filipi & Jack, 2020). In healthy volunteers, IFN-β symptoms peak around 6-8 hr post-injection (Exton et al., 2002; Salmon et al., 1996). Existing models of experimental inflammation, such as typhoid or endotoxin challenges, present limitations. Typhoid challenges are often less effective in eliciting a robust immune response, while endotoxin challenges, although effective, can be highly invasive and pose significant risks to participants.

This study investigates the physiological and behavioural response to peripheral IFN-β administration in healthy young (18-34) and older (60-75) adults, focusing on age-related modulation of these effects. In a randomized, placebo-controlled, repeated measures cross-over design, 30 participants were tested on two separate occasions and received a subcutaneous injection in the abdomen. In one session this was reconstituted IFN-β EXTAVIA® and the other saline (placebo). Physiological measures and self-report behavioural questionnaires were collected hourly for the duration of the testing sessions. Blood samples for cellular and immune response were collected at baseline, 4 and 6 ½ hr post-challenge. Blood samples for whole-blood mRNA analysis were collected at baseline and 6 ½ post injection.

By presenting this new model, this study aims to provide a more ecologically valid alternative for studying the impacts of inflammation, potentially enhancing the understanding and treatment of various neuroinflammatory and neuropsychiatric conditions. Additionally, we will compare and discuss the IFN-β response in relation to other experimental challenges, thereby broadening the scope of insights into immune responses and their implications for these conditions.

## METHODS

### Participants

30 healthy participants (15 young [6 male, mean age 25.2 ± 5.1 years], 15 old [6 male, mean age 65.6 ± 4.5 9 years]), were recruited from around the Cardiff area. Volunteers had to be non-smokers and in good health, as determined by their medical history, physical and psychiatric screening, vital signs and clinical laboratory test results including renal, liver, thyroid function and full blood count. In the younger cohort, the majority of participants identified as belonging to a white ethnic background (n=13), with two participants identifying as of Asian origin. The older cohort was composed of participants who identified as being of white ethnicity. Participants were advised to avoid heavy exercise and the use of alcohol 24 hr before the sessions. The study was approved by the London-Camden & Kings Cross Research Ethics Committee (20/LO/0239)

### Study Design

We adopted a randomised, placebo-controlled, repeated measure cross-over study. All participants were tested on two separate occasions between 2 and 4 weeks apart (M= 28.2 days). In one session this was 4mL of reconstituted IFN-β EXTAVIA® (100 µg), and in the other 4mL of 0.9% saline. The order of intervention was randomised with half of the participants receiving IFN-β on their first study session and half placebo.

Temperature, heart rate, systolic and diastolic blood pressure were measured at baseline and 6 further time-points in both study sessions (1 hr, 2 hr, 3 hr, 4 hr, 5½ and 6½ hour post-IFN-β/saline injection). Self-report mood, fatigue and sickness questionnaires (POMS, fVAS and SicknessQ) were administered at baseline, and 5 additional time points. Blood samples were collected at baseline, 4 hr and 6½ hr post-challenge for haematological analysis and whole-blood mRNA analysis at baseline and 6 ½ hr post-injection

### Cytokine analysis

Blood samples for cytokine analysis were collected into BD Vacutainer plastic EDTA tubes with lavender hemogard closure (4 mL, 13×75mm) (Becton, Dickson and Company, Franklin Lakes, New Jersey, United States) at baseline, 4 hr and 6 ½ hr post-injection. EDTA tubes were centrifuged immediately at 2000 rpm for 20 min. Plasma was removed, aliquoted and stored at -80 °C before analysis. IFN-β plasma concentration was measured using VeriKine-HS™ Human IFN Beta Serum High Sensitivity ELISA Kit (PBL Assay Science, NJ, USA). Detection limit was 2.3 pg/mL and intra and inter-essay coefficients of variation were 3.6% and 7.9%.

Plasma levels of IL-6, TNF- α and IL-10 were quantified with Quantikine™ High Sensitivity (R&D Systems inc., Minneapolis, USA). Detection limits were 0.156 pg/mL, 0.156 pg/mL and 0.78 pg/mL respectively and coefficients of variation were 3.6% and 4.9% (IL-6), 2% and 6.7% (TNF- α) and 5.8% and 7.8% (IL-10). Standards and samples were tested in duplicate. Samples with measurements below the lowest standard were given a value of half the lower limit of detection for IL-10 and IFN-β cytokines (Breen et al., 2011)

### Transcriptomics Analysis

Blood samples were collected in PAxGene RNA tubes and stored at -80 °C. RNA was extracted following the PAXgene Blood RNA Kit, including a globin depletion step. RNA quality and concentration were assessed using Qubit and fragment analysis (https://www.thermofisher.com/uk/en/home/industrial/spectroscopy-elemental-isotope-analysis/molecular-spectroscopy/fluorometers/qubit/qubit-assays.html) before samples were shipped on dry ice to Lexogen for 3′ mRNA sequencing using the QuantSeq protocol (https://www.lexogen.com/quantseq-family/).

Differentially expressed genes (DEGs) were analysed through the use of IPA (QIAGEN Inc.,https://www.qiagenbioinformatics.com/products/ingenuity-pathway-analysis). To identify significant pathways, IPA uses the p-value of overlap, calculated using the right-tailed Fisher’s exact Test. Differentially expressed genes were identified (|log2FC| > 1 and adjusted p-value < 0.05) using edgeR/limma (https://ucdavis-bioinformatics-training.github.io/2022-April-GGI-DE-R/data_analysis/DE_Analysis_with_quizzes_fixed). Volcano plots were generated using the EnhancedVolcano R package and R.4.4.2.

### Statistical Analysis

The main effects of INF-β and age, as well as their interaction, on physiological, cellular, immune, and behavioural data, were analysed using repeated measures mixed factorial ANOVAs: within-subject factors: condition (IFN-β/placebo) and time (pre- and post-injection times as described), between-subject factor: age (young/old) with significant interactions assessed using t-tests at each time points between conditions. Pearson’s correlations were used to assess associations between cytokines and total and differential cell counts (computed as peak change minus baseline for the IFN-β condition). Data analysis was carried out using SPSS 27 statistical package.

## RESULTS

### PHYSIOLOGICAL RESPONSE

#### Temperature

A repeated measures ANOVA revealed a significant main effect of condition (placebo/IFN-β) (F(1,28) = 45.5, p < 0.001) (+1.1 °C in the IFN-β condition) and a significant condition x time interaction (F(3.3,94.48) = 30.88, p < 0.001). IFN-β significantly increased temperature compared to placebo from 3 hr post-injection (t(28) = -2.87, p = 0.008), peaking at 6 ½ hr post-injection (t(28) = -8.32, p < 0.001). A significant main effect of age was observed (F(1,28) = 6.79, p = 0.014), with older individuals showing a mean temperature decrease of 0.271°C (SE = 0.104) compared to younger participants, regardless of condition or time (*Fig 1*). No significant condition x age interactions were found.

**Fig 1.**
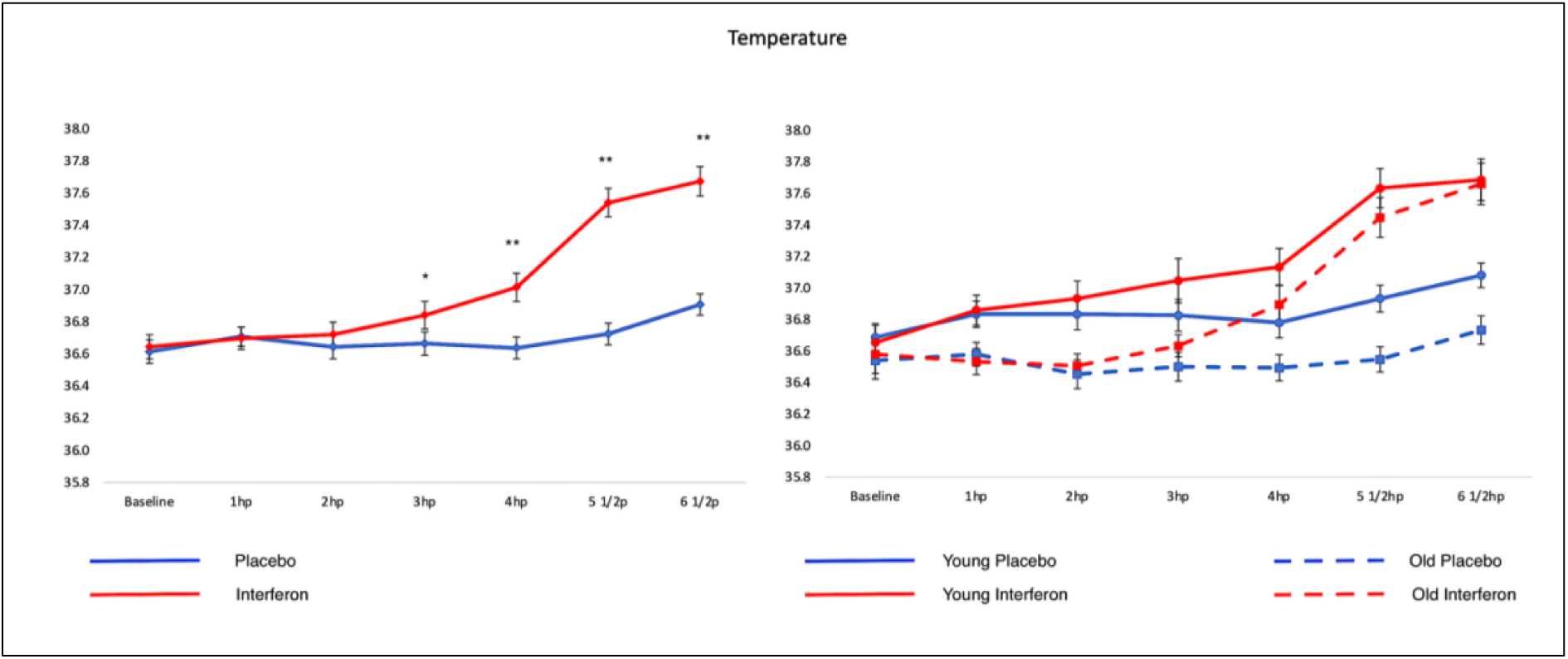
Effects of IFN- β on Temperature. Effects of IFN-β on temperature (°C) (left), and age-associated effects on temperature (right). Red lines represent IFN-β and blue lines placebo. Dotted lines represent older individuals. Error bars denote SEM; significant values show the main effect of IFN-β compared to placebo (*p<0.05 **p<0.001).

#### Heart Rate

The ANOVA indicated a significant main effect of condition on heart rate (F(1,28) = 21.34, p < 0.001) (+11 bpm in the IFN-β condition) and a significant condition x time interaction (F(4.14,115.93) = 16.62, p < 0.001). IFN-β increased heart rate relative to placebo from 3 hr post-challenge (t(28) = -5.6, p = 0.005), peaking at 6 ½ hr post-injection (t(28) = -10.33, p < 0.001) (Fig 2). There were no significant condition x age interactions. However, a significant main effect of age on heart rate was found (F(1,28) = 6.27, p = 0.018), with older individuals having a lower resting heart rate than younger participants (mean difference = 7.657 bpm, SE = 3.05) (*Fig 2*).

**Fig 2.**
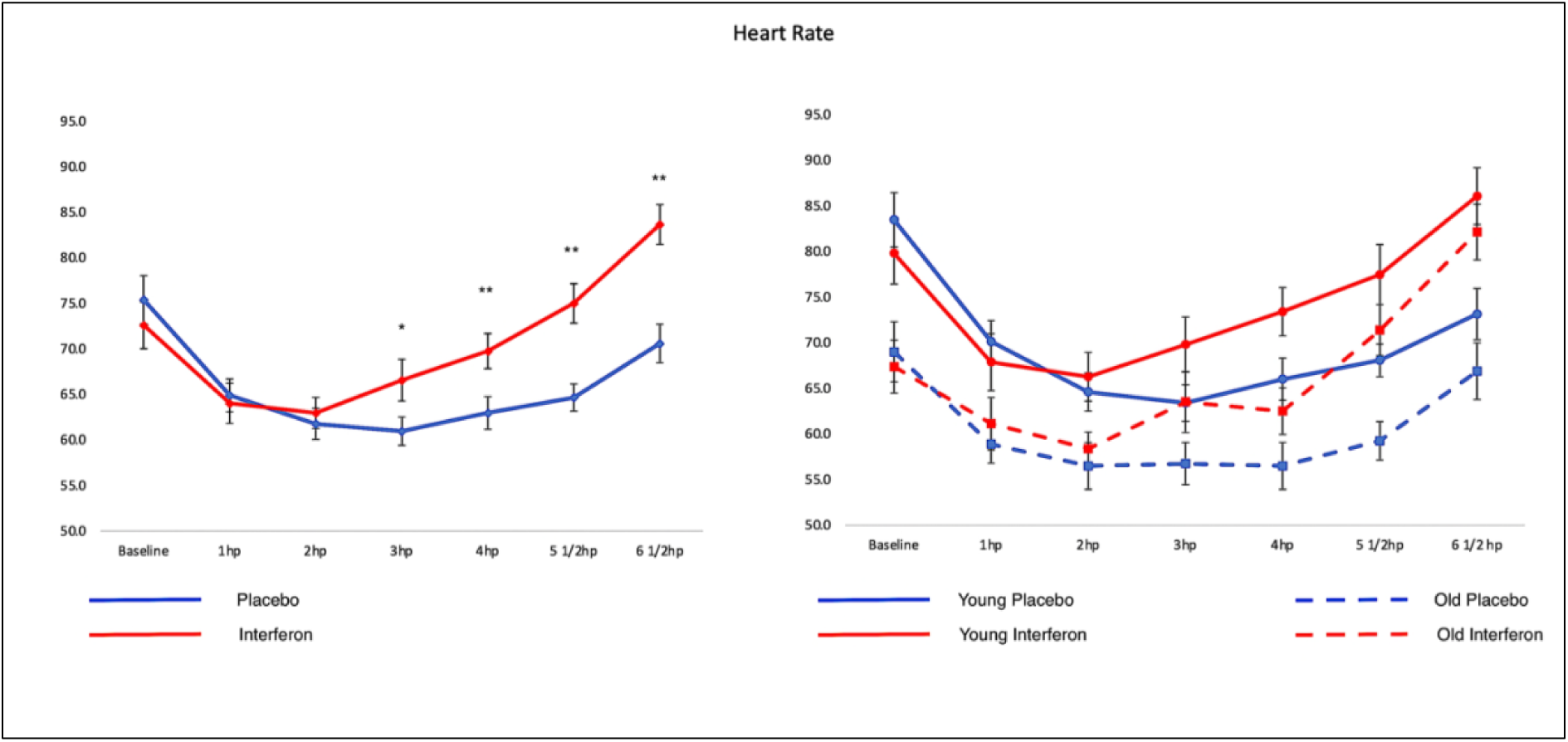
Effects of IFN- β on heart rate. Effect of IFN-β on heart rate (bpm)(left) and age-associated differences on heart rate (right). Red lines represent IFN-β and blue lines placebo. Dotted lines represent older individuals. Error bars denote SEM; significant values show the main effect of IFN-β compared to placebo (*p<0.01 **p<0.001).

#### Blood Pressure

For systolic blood pressure, repeated measures ANOVAs showed a marginal main effect of condition (F(1,28) = 4.15, p = 0.051) but no significant condition x time or age interactions. No significant differences were observed in diastolic blood pressure or its interactions (*Fig 3*). Between-subject effects revealed a significant age-associated difference in systolic BP (F(1,28) = 14.58, p < 0.001), with older individuals exhibiting higher systolic BP compared to younger participants (mean difference = -19.086 mmHg, SE = 4.9) (*Fig 3*). No age-associated effects were found for diastolic BP.

**Fig 3.**
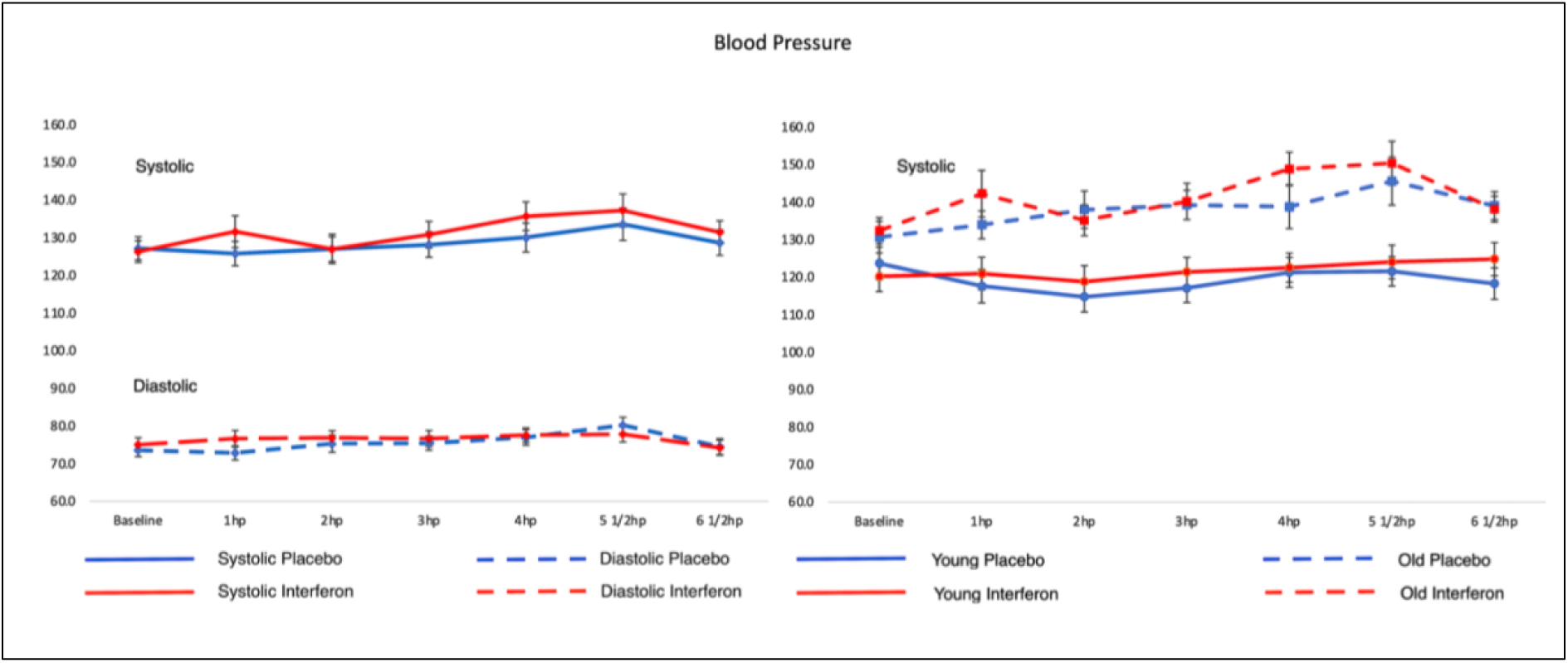
Effects of IFN- β on blood pressure. Effects of IFN-β on Blood pressure (mmHg) (left) and age-associated effect of systolic BP (right). Left: Red lines represent IFN-β, blue lines placebo, dotted lines represent diastolic BP. Right Fig: dotted lines represent older individuals. Error bars denote SEM.

## CELLULAR IMMUNE RESPONSE

Significant condition x time interactions were found for total WBC (F(1.49,40.47) = 10.92, p < 0.001) and differential counts: monocytes (F(1.52,42.79) = 7.22, p < 0.001), neutrophils (F(1.52,42.43) = 15.9, p < 0.001), and lymphocytes (F(1.33,37.32) = 72.46, p < 0.001). A main effect of condition was observed for lymphocytes (F(1,28) = 112.26, p < 0.001) and the neutrophil to lymphocyte ratio (NLR) (F(1,28) = 35.95, p < 0.001), with a significant condition x time interaction for NLR (F(1.06,29.7) = 45.78, p < 0.001) (*Fig 4*).

**Fig 4.**
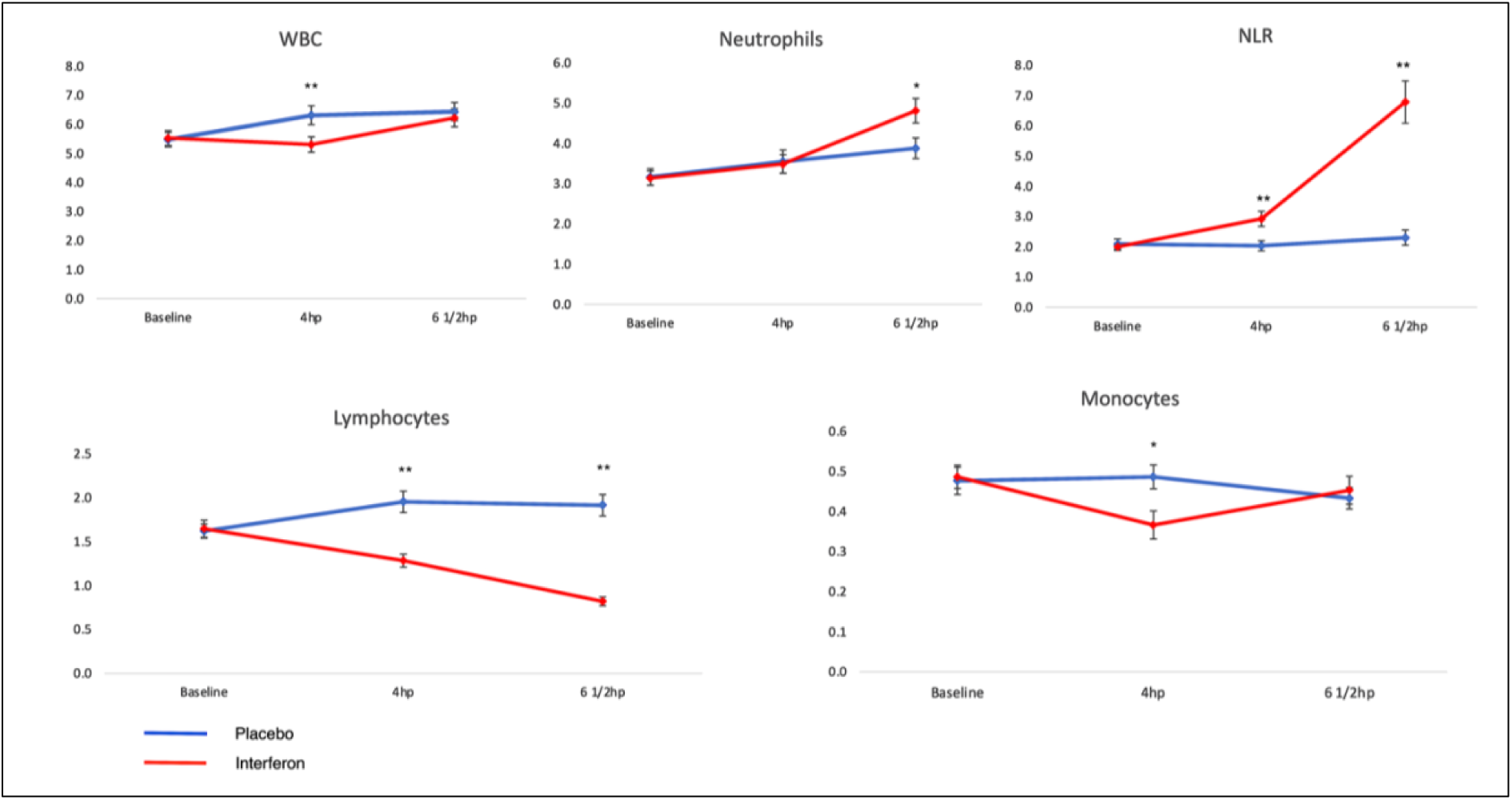
Effects of IFN- β on total and differential WBC and Neutrophil to Lymphocyte ratio. Effects of IFN-β on total and differential WBC and Neutrophil to Lymphocyte ratio (NLR) (10^9^/L). Red lines represent IFN-β, blue lines placebo. Error bars denote SEM; significant values show the main effect of IFN-β relative to placebo (*p<0.01,**p<0.001).

Post-IFN-β, cell counts (expressed as relative delta percentage ± CI) increased for neutrophils (59.2 ± 20.4%) at 6 ½ hr post-injection. NLR increased at both 4 hr (43.8 ± 11.6%) and 6 ½ hr (245.4 ± 67.2%) post-IFN-β. Lymphocytes decreased by 21.2 ± 4.8% and 49.3 ± 4.8% at 4 and 6 ½ hr, respectively. Monocytes decreased by 29.6 ± 8.5% at 4 hr post-IFN-β, with WBC also showing a decrease of 3.5 ± 5.8% at 4 hr relative to baseline.

A significant between-subject effect was found for monocytes (F(1,28) = 7.93, p = 0.026), with older individuals showing lower monocyte counts regardless of time and condition (mean difference = 0.126 (10^9^/L), SE = 0.045). A non-significant trend for condition x age interaction was observed for lymphocytes (F(1,28) = 3.13, p = 0.088) (*Fig 5*). No other interactions or age-associated differences were observed for other WBC counts. Haematology data are presented in Table 1.

**Fig 5.**
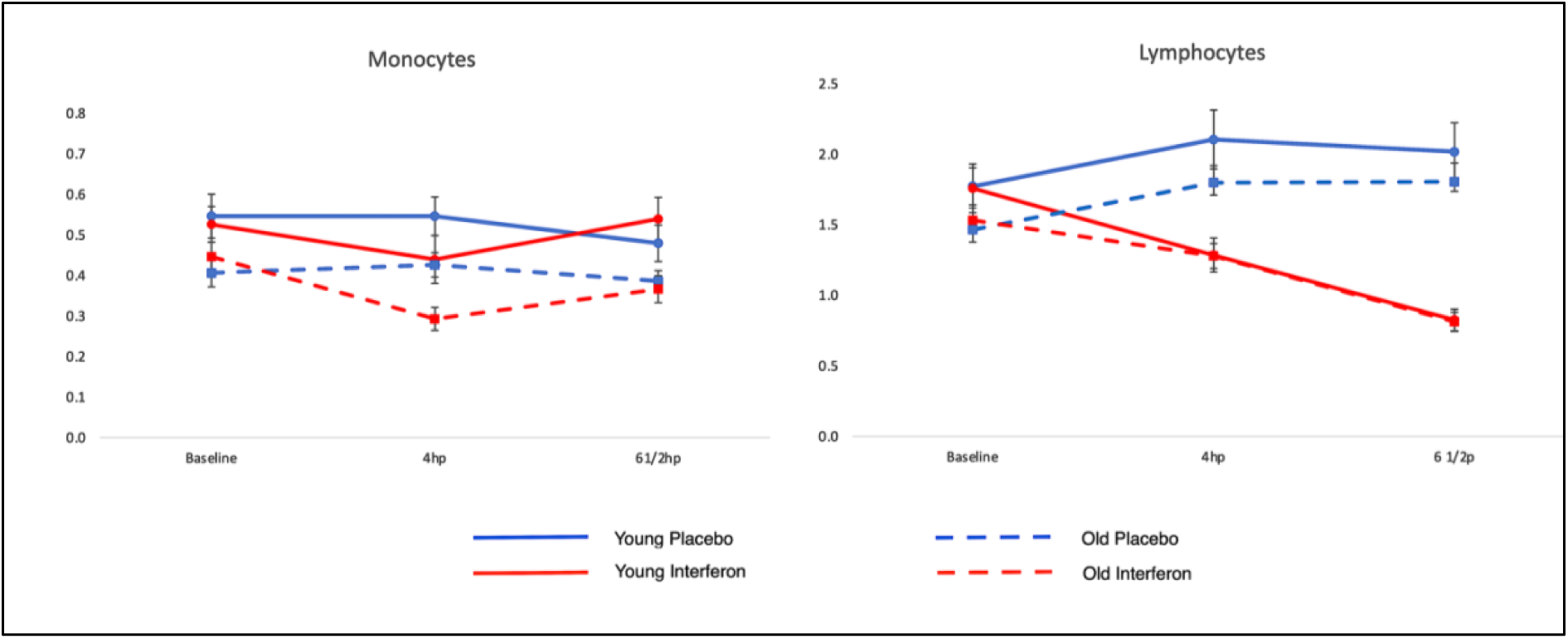
Age-associated effects on Monocytes and condition x age interaction on Lymphocytes. Age-associated effects on Monocytes (left) and condition x age interaction on Lymphocytes (right) (10^9^/L). Red lines represent IFN-β, blue lines placebo. Dotted lines represent older participants. Error bars denote SEM.

**Table 1.** Haematology data ([10^9^/L] ± SEM)

| | Time post-injection (hrs) | Saline | IFN-β | p-values (Saline vs. IFN-β) |
| --- | --- | --- | --- | --- |
| Total WBC | 0 | 5.48 (0.24) | 5.53 (0.25) | .789 |
|  | 4 | 6.32 (0.32) | 5.31 (0.26) | <.001 |
|  | 6 1/2 | 6.44 (0.31) | 6.23 (0.31) | .478 |
| Monocytes | 0 | 0.47 (0.03) | 0.48 (0.02) | .759 |
|  | 4 | 0.48 (0.02) | 0.34 (0.02) | <.001 |
|  | 6 1/2 | 0.43 (0.02) | 0.42 (0.02) | .849 |
| Lymphocytes | 0 | 1.62 (0.08) | 1.64 (0.09) | .699 |
|  | 4 | 1.95 (0.12) | 1.28 (0.07) | <.001 |
|  | 6 1/2 | 1.91 (0.12) | 0.82 (0.05) | <.001 |
| Neutrophils | 0 | 3.17 (0.2) | 3.14 (0.170) | .848 |
|  | 4 | 3.67 (0.25) | 3.49 (0.23) | .414 |
|  | 6 ½ | 3.88<br>(0.25) | 4.81<br>(0.3) | .003 |
| NLR | 0 | 2.08<br>(0.16) | 2.0<br>(0.11) | .536 |
|  | 4 | 2.03<br>(0.16) | 2.92<br>(0.24) | <.001 |
|  | 6 ½ | 2.3<br>(0.25) | 6.79<br>(0.69) | <.001 |

## CYTOKINE RESPONSE

Following the IFN-β challenge, plasma concentrations of IFN-β (pg/mL) significantly increased over time (F(1.19,32.19) = 10.14, p < 0.001). Compared to baseline, concentrations peaked at 4 hr (t(29) = -8.74, p < 0.001) and decreased at 6 ½ hr post-challenge (t(29) = -8.55, p < 0.001), with significant differences between the 4 and 6 ½ - hour marks (t(29) = 6.98, p < 0.001). From baseline, INF- β levels increased approximately 15-fold at 4 hr and 9-fold at 6 ½ hr. Although no significant age effects were observed, there was a trend towards a time x age interaction, with higher IFN-β levels observed in younger individuals (F(1.18,33) = 2.8, p = 0.098) (*Fig 6*).

**Fig 6.**
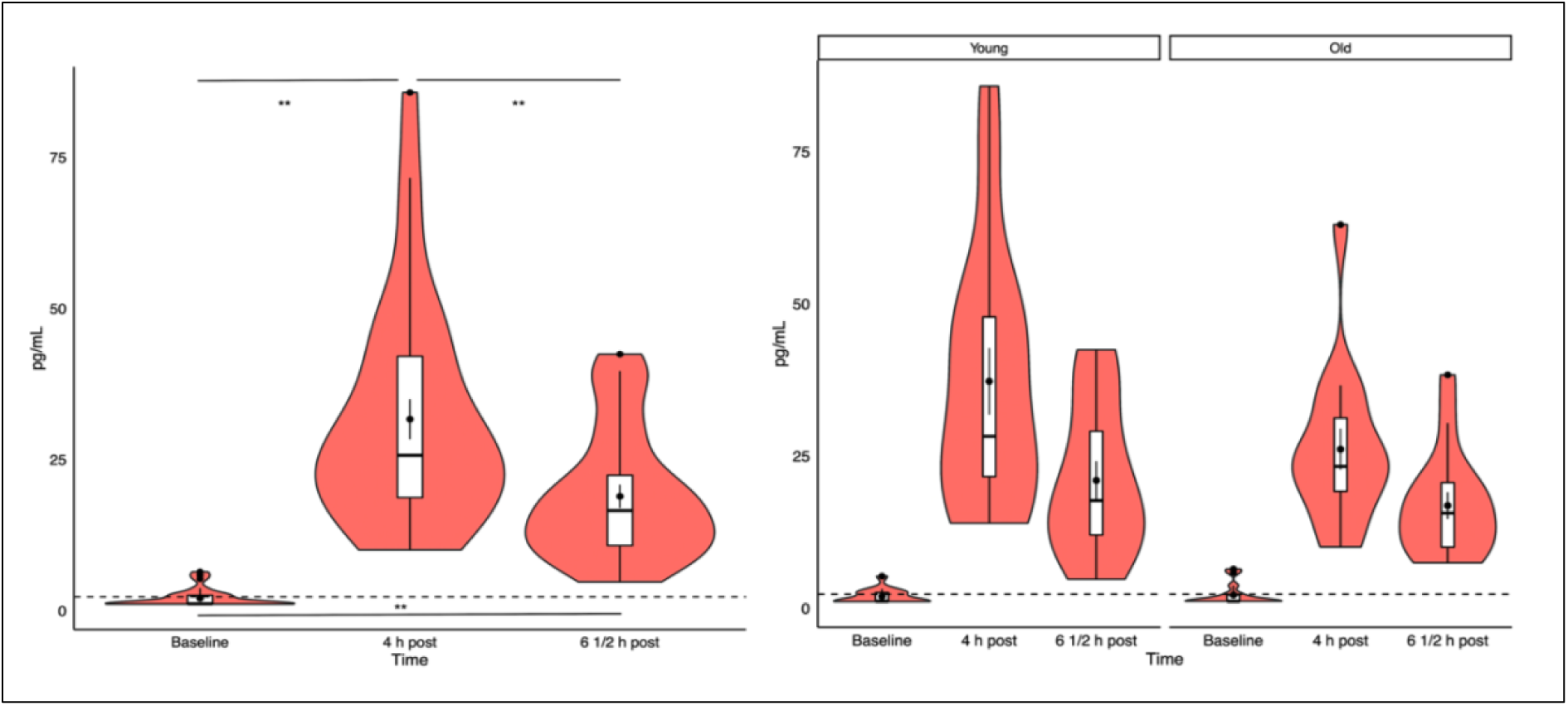
Distribution of IFN- β plasma concentrations for the interferon condition. Distribution of IFN-β plasma concentrations (left) and split by age (right) for the interferon condition. Significant values show main paired sample t-test results (**p<0.001). Dashed line shows lower limit of detection (2.3 pg/mL).

IFN-β also significantly increased circulating IL-6 levels (approximately 6-fold), as shown by significant condition effects (F(1.28) = 10.14, p = 0.004) and condition x time interactions (F(1.45,40.78) = 4.49, p = 0.027). A time x age interaction was noted (F(1.85,51.83) = 4.16, p = 0.024), with mean ± SEM IL-6 concentrations of 6.98 ± 1.1 (young) and 8.25 ± 1.1 (old) at 4 ½ hr, and 8.38 ± 1.4 (young) and 13.24 ± 1.4 (old) at 6 ½ hr post-injection. Additionally, there was a trend towards a condition x time x age interaction, with older individuals showing higher IL-6 concentrations at 6 ½ hr post-injection (F(1.45,40.78) = 2.65, p = 0.097) *(Fig 7*).

**Fig 7.**
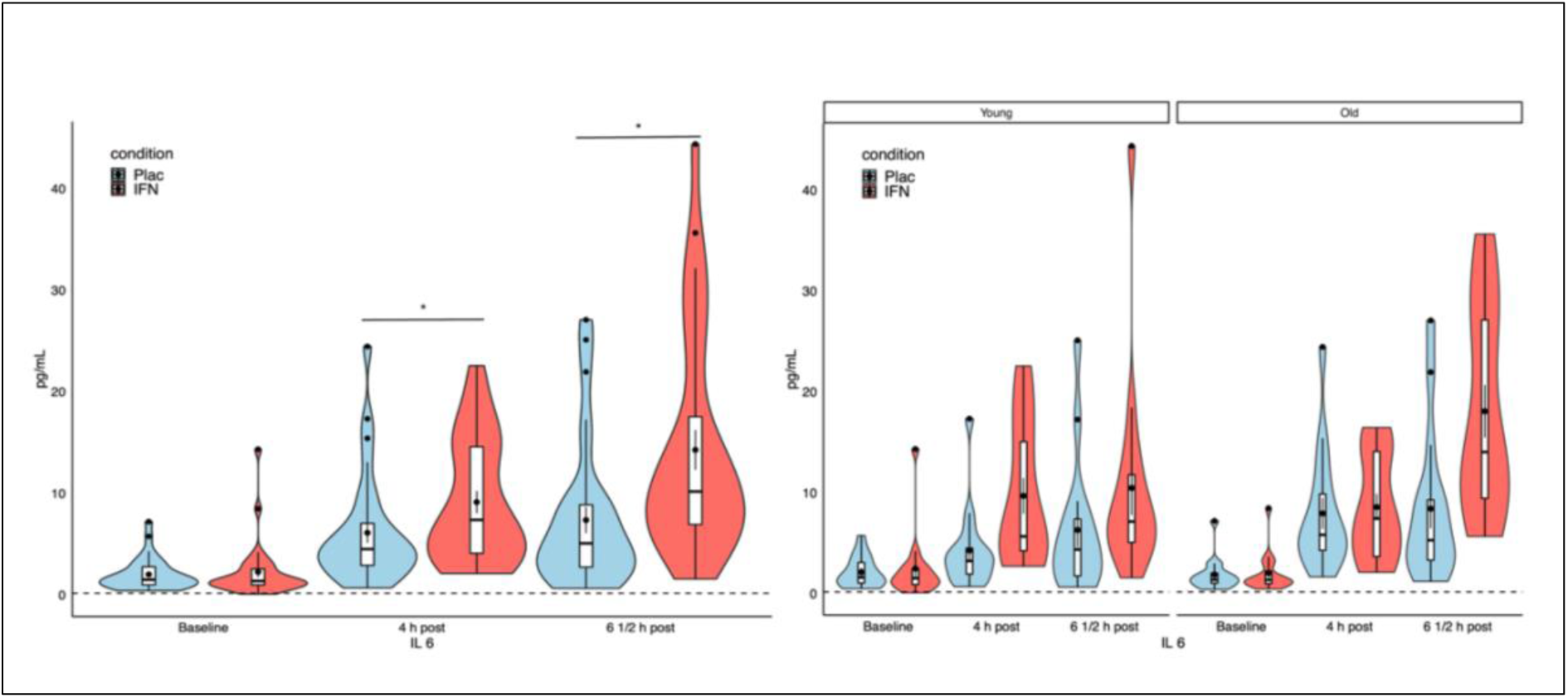
Distribution of IL-6 plasma concentrations. Distribution of IL-6 plasma concentrations (left) and IL-6 plasma concentrations split by age (right). Blue plots denote placebo, red plots interferon. Significant values show paired sample t-test results (*p<0.05). Dashed line shows lower limit of detection (0.156 pg/mL).

Condition x time interaction effects were observed for TNF-α (F(1.56,43.94) = 22.07, p < 0.001) with an average increase of approximately 1.3 fold following IFN- β (*Fig 8).* No condition x age interactions or age-associated differences were found. There was no significant main effect of IFN-β on IL-10 concentrations (p > 0.1). However, between-subjects effects revealed a significant age-associated difference (F(1.28) = 6.48, p = 0.017), with older individuals showing lower IL-10 levels compared to younger participants (mean difference = 0.889 pg/mL, SE = 0.349) (*Fig 9*). Cytokine data and correlations are detailed in Tables 2 and 3, respectively.

**Fig 8.**
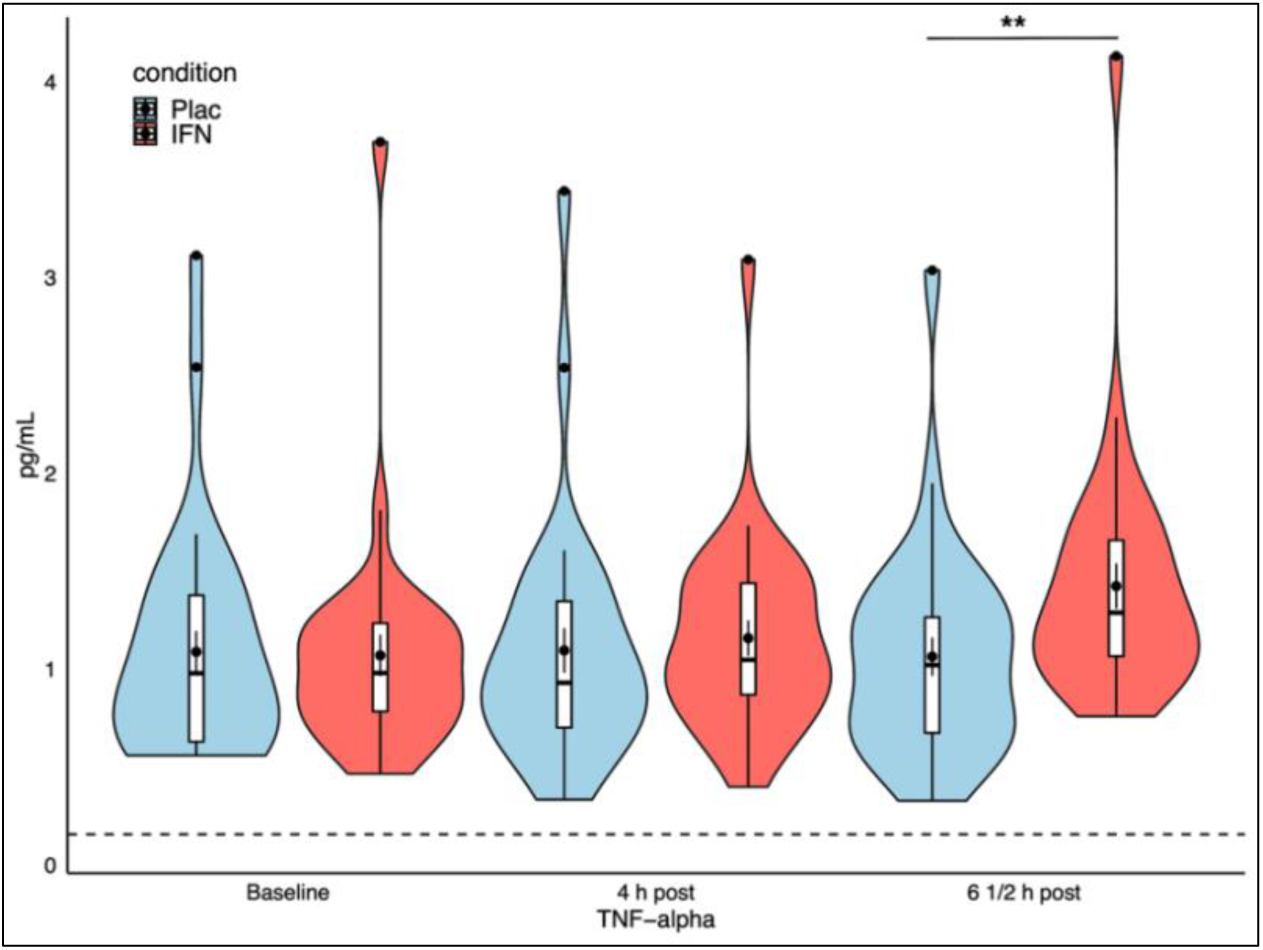
Distribution of TNF- α plasma concentrations. Red bars represent IFN-β, blue placebo Significant values show paired sample t-test results (**p<0.001). Dashed lines show lower limit of detection (0.156 pg/mL).

**Fig 9.**
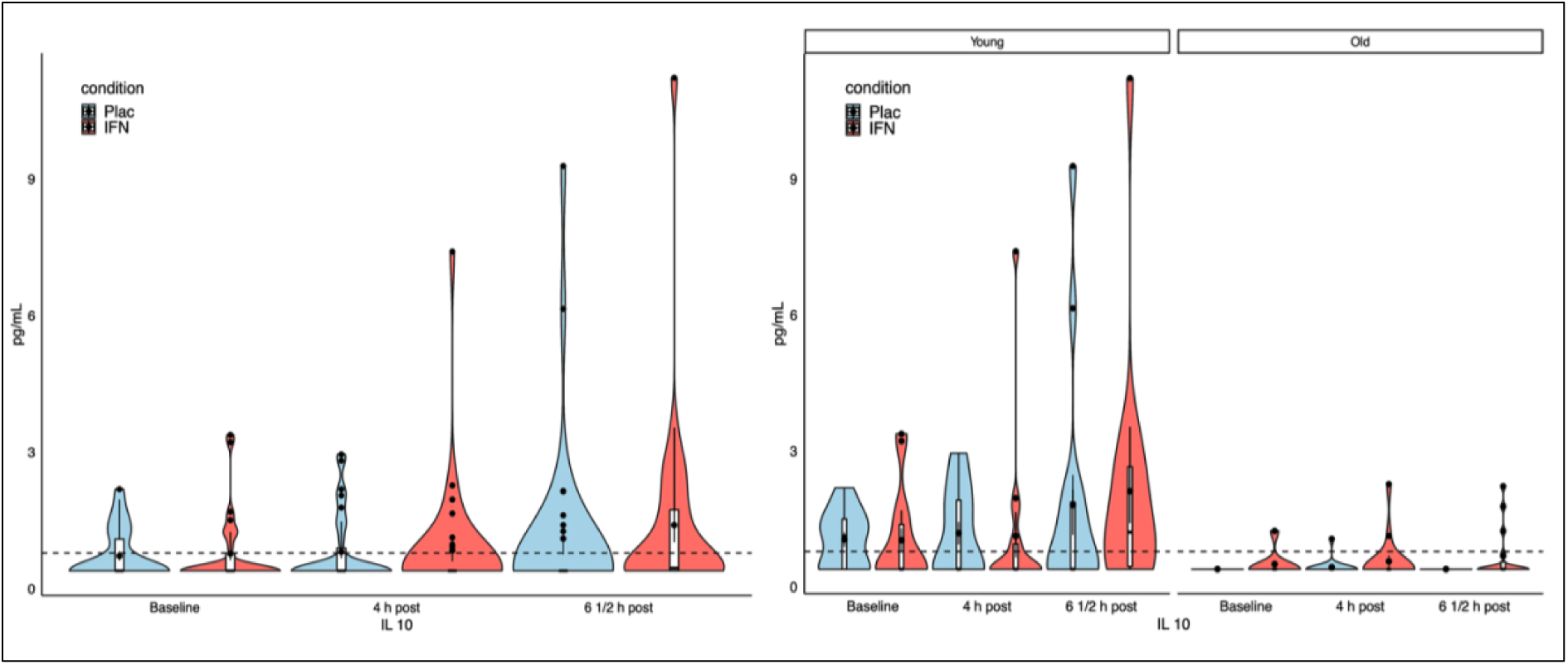
Distribution of IL-10 plasma concentrations. Distribution of IL-10 plasma concentrations (left) and IL-10 plasma concentrations split by age (right). Blue plots denote placebo, red plots interferon. Dashed line shows lower limit of detection (0.78 pg/mL).

**Table 2.** Cytokine data ([pg/mL] ± SEM)

| IFN-β | 0 | 4 | 6 ½ | p-values<br>(relative to<br>baseline) |  |
| --- | --- | --- | --- | --- | --- |
|  |  |  |  | (4 hr) | (6 ½<br>hr) |
|  | 2.05<br>(0.27) | 31.69<br>(3.32) | 18.92<br>(1.92) | <.001 | <.001 |
|  | Time<br>post-injection<br>(hrs) | Saline | IFN-β | p-values<br>(Saline vs. IFN-β) |  |
| IL-6 | 0 | 2.0<br>(0.28) | 2.24<br>(0.51) | .609 |  |
|  | 4 | 6.1<br>(0.98) | 9.13<br>(1.09) | .036 |  |
|  | 6 ½ | 7.35<br>(1.29) | 14.26<br>(1.96) | .011 |  |
| IL-10 | 0 | 0.71<br>(0.1) | 0.76<br>(0.14) | .744 |  |
|  | 4 | 0.8<br>(0.14) | 0.84<br>(0.24) | .891 |  |
|  | 6 ½ | 1.09<br>(0.35) | 1.39<br>(0.37) | .225 |  |
| TNF- α | 0 | 1.08<br>(0.1) | 1.06<br>(0.1) | .828 |  |
|  | 4 | 1.09<br>(0.11) | 1.15<br>(0.09) | .512 |  |
|  | 6 ½ | 1.06<br>(0.98) | 1.42<br>(0.11) | <.001 |  |

**Table 3.**
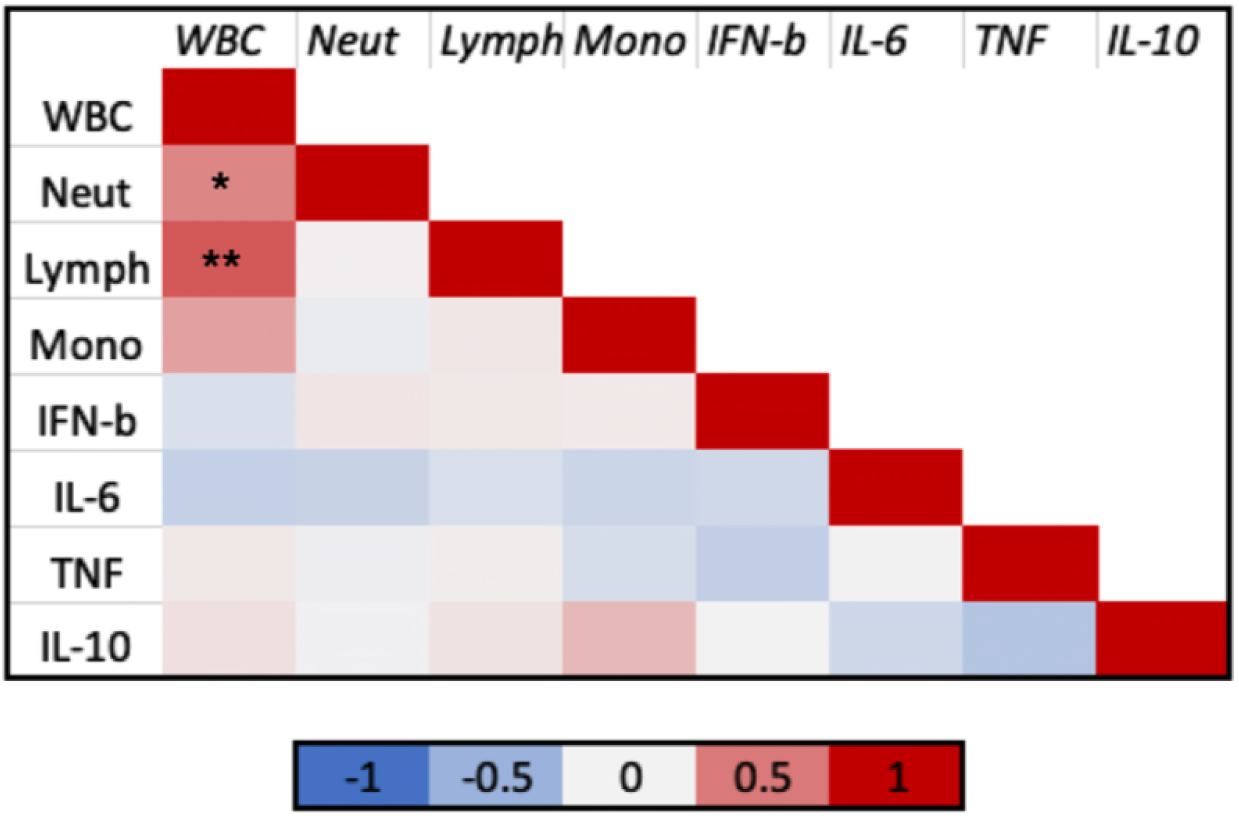
Cytokine correlations with immune cells.

## TRANSCRIPTOMIC RESPONSE

Total RNA from the blood of the subjects treated with IFN-β was analysed using RNA-sequencing to quantify differentially expressed genes (DEGs) relative to the placebo control group. DEGs were analysed through the use of IPA (QIAGEN). To identify significant pathways, IPA uses the p-value of overlap calculated using the right tailed Fisher’s exact Test. As expected, IFN-β treatment triggered IFN-α/β signaling pathways, as well as innate immune pathways leading to inflammation, such as the Pathogen-induced cytokine storm signalling pathways, pattern recognition pathways, inflammasome, Toll-like receptor signalling pathways, cGAS-STING as well as RIG-I and TLR3 pathways (*Fig* 10A). The overview of the main biological pathways (*Fig* 10B) demonstrates how these pathways relate to one another (Fig 10B) and most importantly how the interferon signalling pathway that is activated is linked to the anti-viral immune response as well as the innate immune pathways that lead to inflammation.

**Fig. 10.**
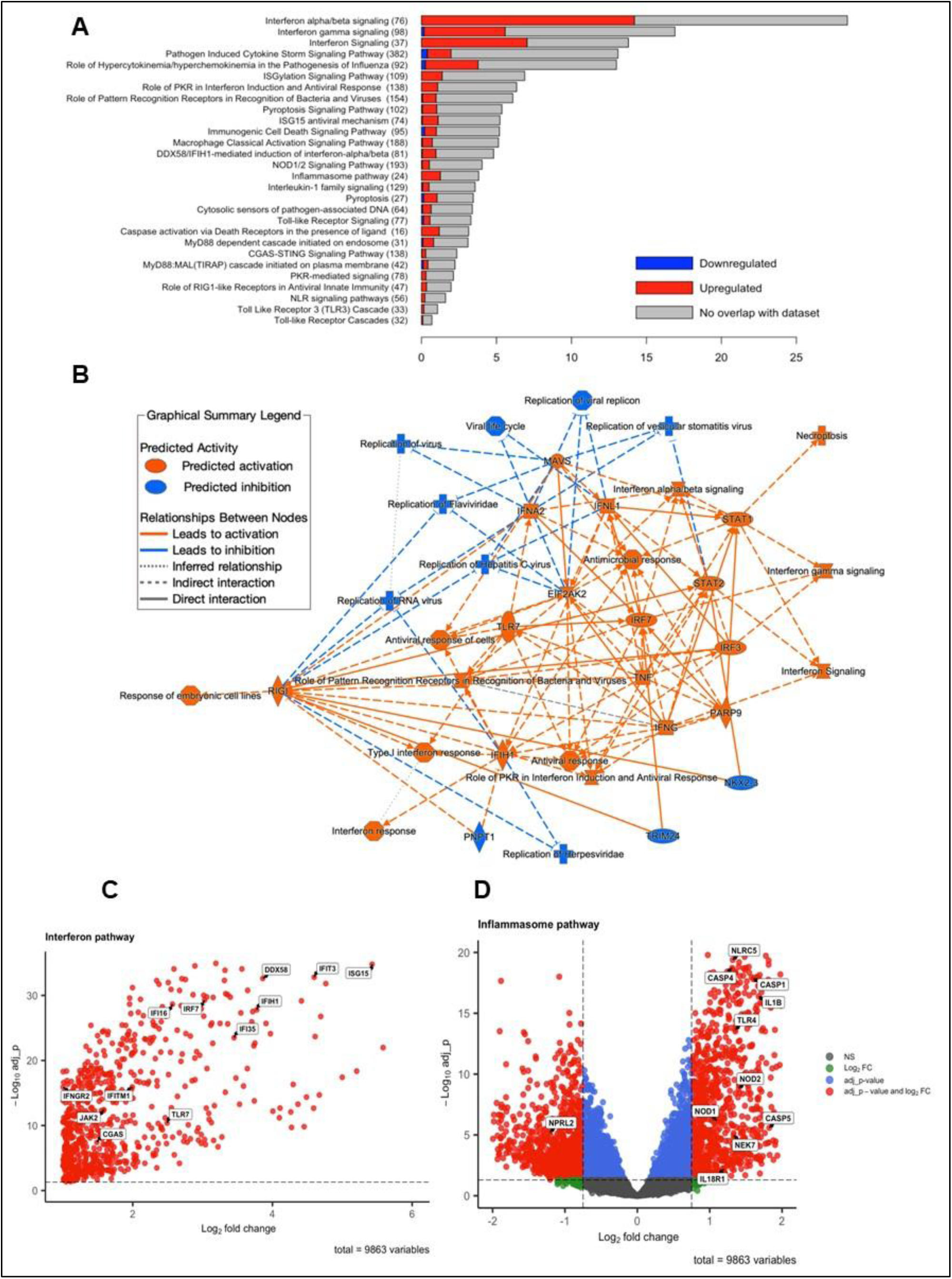
Transcriptomics analysis of RNA-sequencing profiling of subjects administered with IFN-β compared to placebo controls. Differentially expressed genes (DEGs) were analysed through the use of IPA. (A) Stacked bar chart displaying the proportion of positive (red) and negative (blue) DEGs relative to the total number of molecules curated for each canonical pathway. The number in parentheses indicates the total number of genes in the canonical pathway. (B) Overview of the main biological entities (canonical pathways, upstream regulators, target molecules) and how these concepts relate to one another. The types of relationship lines are summarized in the Graphical Summary Legend. Differentially expressed genes were identified (|log2FC| > 1 and adjusted p-value < 0.05) using edgeR/limma. Volcano plots summarising significant genes are displayed (C,D). Highlighted are key DEGs involved in the interferon (C) and inflammasome pathways (D).

Analysis of DEGs identified 1239 differentially expressed genes and showed upregulation of genes involved in the interferon pathways such as ISG15, IFIT3, IFIH1, IFI35, DDX58, IRF7, IFI16, IFITM1, and IFNGR2 (*Fig* 10C). Cytosolic nucleic acid sensors, such as TLR7 and cGAS, which once activated trigger IFN-β, were some of the genes that were upregulated (*Fig* 10C). In addition, several genes of the inflammasome pathway were found to be upregulated such as IL1B, IL18R1, NOD1, NOD2, NLRC4 and NEK7, which regulate the NLRP3 inflammasome (*Fig* 10D) (He et al., 2016). NEK7 is an essential mediator of NLRP3 activation downstream of potassium efflux. Interestingly, there also was also upregulation of CASP1, CASP4 and CASP5 genes suggesting that there is canonical (CASP1) as well as non-canonical (CASP4, CASP5) inflammasome activation. The results indicate for the first time that administration of IFN-β as an in-vivo experimental-medicine model of human inflammation is able to trigger a robust IFN-related and inflammasome-related gene signature similar to a response to infection.

## SUBJECTIVE SICKNESS RESPONSE

Interferon-beta induced significant mood shifts observed in five POMS subscales, with the most pronounced changes at 5.5 hr post-injection. IFN-β significantly decreased total mood score (Condition x (Placebo/IFN-β) x Time F(3.13,87.82) = 5.93, p < 0.001) and vigour (F(3.5,98.58) = 3.19, p = 0.021), and increased negative mood (F(3.28,91.98) = 5.61, p < 0.001), tiredness (F(2.44,68.5) = 4.97, p = 0.006), and tension (F(3.89,109.13) = 2.73, p = 0.034).

Sickness symptoms increased significantly, as shown by the SicknessQ, with a significant condition effect (F(1,28) = 7.64, p = 0.01) and condition x time interaction (F(2.88,80.87) = 7.39, p = 0.001), peaking at 5.5 hr post-injection (*Fig 11*). No significant effects were found in the fatigue scale (fVAS), and no condition x age interactions or age-related effects were observed for any scales (p>0.1).

**Fig 11.**
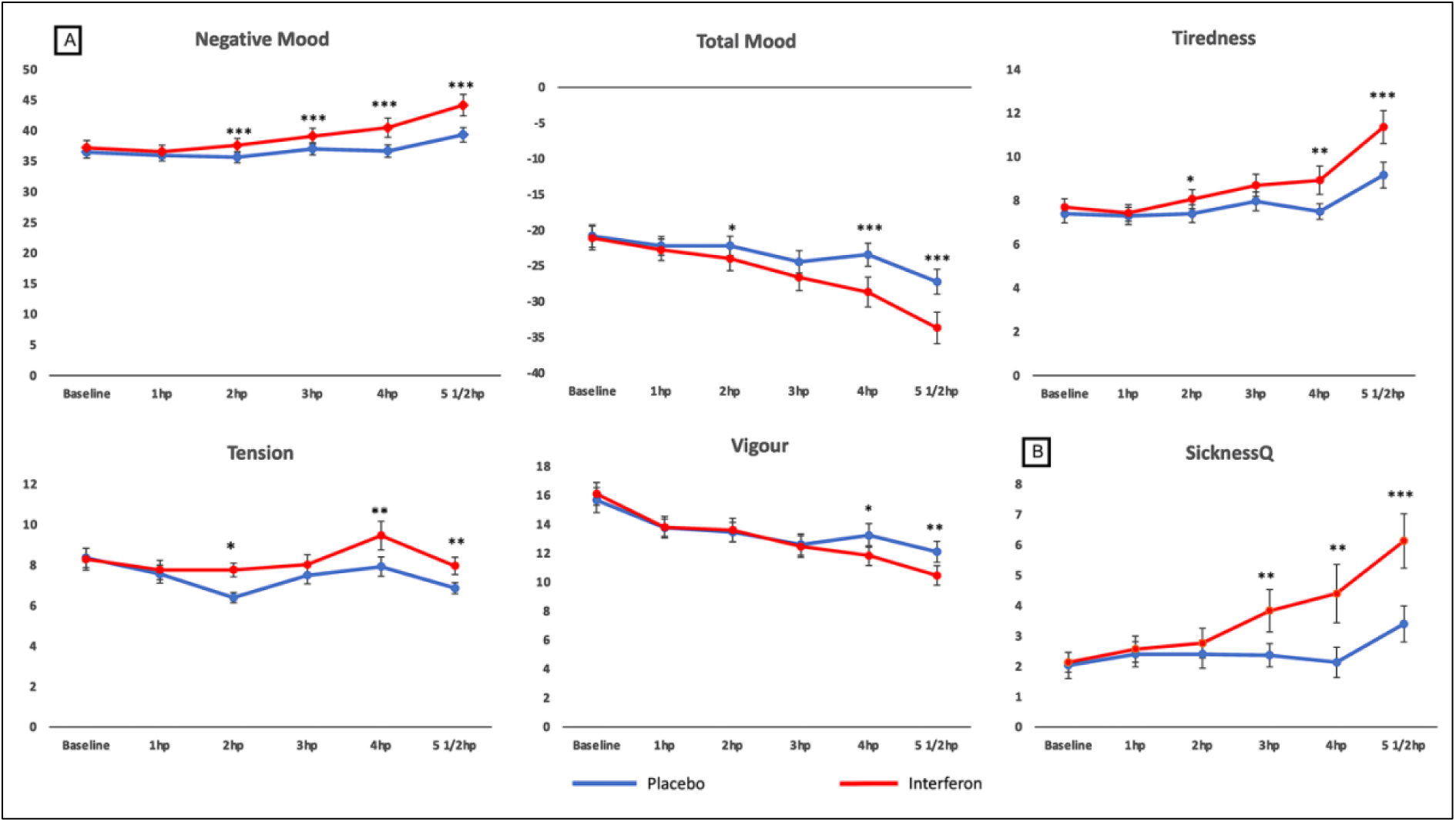
Subjective response. (A) Significant POMS subscales; (B) Sickness Questionnaire (SicknessQ). Blue represents placebo, red interferon. Error bars denote SEM. Significant values show the main effect of IFN-β relative to placebo (* p< 0.05, ** p< 0.01, *** p< 0.001).

### Side effects

Routes of administration and frequency of injection vary among treatment strategies, nonetheless, in all cases, the IFN injection elicits a systemic inflammatory response that mimics a viral infection and commonly induces a physiological response marked by a variety of flu-like symptoms. (Davis et al., 1989; Filipi & Jack, 2020). In healthy individuals, IFN-β has shown a half-life and bioavailability of approximately 4-5 hr and 30-50% respectively, reaching peak concentrations between 1-8 hr post-injection. Common clinical side effects, such as chills, fever, fatigue, nausea, are generally mild, appear approximately 3–4 hr post-infusion, and are consistent across different administration routes (Hu et al., 2016; Salmon et al., 1996). Injection site reactions are also common; however, in our study, only one participant experienced mild itchiness and redness in the injection area, which resolved within 24 hr. This reaction was likely due to the reconstituted interferon solution being refrigerated briefly prior to administration rather than injected immediately after preparation. Overall, symptoms were mild and resolved by the evening of the intervention for most participants. Most older individuals reported experiencing more pronounced symptoms, with some persisting overnight and into the following morning. These observations were based on participant feedback collected during informal post-session discussions.

## DISCUSSION

In this study, we establish IFN-β as a safe and reliable experimental model of acute inflammation. Our findings demonstrate that IFN-β elicits a robust systemic immune response, characterised by changes in physiological and behavioural measures. This includes elevated temperature and heart rate, immune cell activation (lymphocytes, monocytes and neutrophils), and heightened cytokine levels (IFN-β, IL-10 and TNF-α). These physiological changes are accompanied by pronounced subjective responses, such as increased negative mood, fatigue and sickness symptoms.

By inducing a transient systemic immune response, IFN-β offers a valuable model for investigating neuroinflammation in healthy individuals. This paper also compares IFN-β induced responses to other experimental inflammatory challenges, including endotoxin (LPS), IFN-α, and typhoid vaccination, highlighting its unique profile. Furthermore, we explore age-related differences in response to IFN-β, paving the way for future research to refine and validate this model for diverse applications.

IFN-β induced a significant systemic inflammatory response, with notable increases in body temperature and heart rate peaking at 6 ½ hr post-injection, consistent with previous studies (Exton et al., 2002; Kümpfel et al., 2000; Salmon et al., 1996). The transient changes in temperature (+1.1 °C) and heart rate (+11 bpm) were similar to those observed with IFN-α but milder compared to LPS and stronger than typhoid models (Fukuhara et al., 1999; Glue et al., 2000; Han et al., 2013; Harrison et al., 2009, 2014; Hijma et al., 2020; Lasselin et al., 2017). IFN-β also significantly increased circulating IL-6 by approximately sixfold, aligning with patterns seen in Hepatitis-C treatments (Capuron & Miller, 2004; Davi es et al., 2020) LPS (Lasselin et al., 2017; Peters van Ton et al., 2021; Sandiego et al., 2015) and typhoid vaccination studies (Harrison et al., 2014, 2016).

IFN-β raised TNF-α levels by approximately 1.3-fold, a response seen in endotoxin studies but not typically in typhoid vaccine or IFN-α models (Davies et al., 2020; Harrison et al., 2009). Nevertheless, acutely, IFN- β has been shown to contribute to the inflammatory response by increasing TNF-α and IL-6 in MS patients (Kümpfel et al., 2000). Although both IFN-α and IFN-β are Type I interferons, their distinct binding affinities and subsequent effects on cellular signalling pathways may explain their differing impacts on TNF-α production (de Weerd et al., 2013; Ivashkiv & Donlin, 2014).

IL-10 levels, which increase in response to IFN-α and LPS, did not significantly change after IFN-β in our study. While IFN-β is known to boost IL-10 in MS, contributing to its anti-inflammatory effects (Kvarnström et al., 2013; Özenci et al., 2000), the lack of a significant IL-10 response here may be due to the lower IFN-β dose used (100 µg) compared to studies with higher doses (250 µg), where IL-10 increases were noted between 6 and 12 hr post-injection and peaked later (Williams & Witt, 1998).

The cellular response to IFN-β was similar to IFN-α, with notable increases in Neutrophil counts (∼50%) and decreases in Lymphocytes (∼50%), while Monocytes initially decreased (∼30%) then increased (∼35%) at 6 ½ hr. These effects were milder compared to the Endotoxin model but followed a similar trend. Overall, IFN-β induced a transient inflammatory state with consistent symptom development, comparable to IFN-α but less intense than Endotoxin, and more pronounced than the Typhoid model (see Table 4).

**Table 4.** Comparison of the IFN-β model with typhoid, IFN-a, and Endotoxin challenges for the physiological and immune response.

| Human systemic inflammation models |  |  |  |  |
| --- | --- | --- | --- | --- |
|  | Typhoid Vaccination (Typhim) | IFN-α2a (3MU) | Our model:<br>IFN-b-1b (EXTAVIA® 100ug) | Endotoxin (1ng/kg NIH reference) |
| Physiological | Temperature: No change | Temperature: + (~0.8 °C) | Temperature: + (~1.1 °C) | Temperature: ++ (~1.4 °C) |
|  | Heart Rate: + (~1-2 bmp) | Heart Rate: ++ (5~6 bmp) | Heart Rate: ++ (~11 bmp) | Heart Rate: +++ (~25 bmp) |
|  | Blood Pressure: dBP +4 mmHg | Blood Pressure : +/- | Blood Pressure : +/- | Blood Pressure : +/- |
|  | Heart rate variability: +LF/HF | Heart rate variability: +LF/HF | *Heart rate variability: expected +LF/HF | Heart rate variability: ++LF/HF |
| Immune/Inflammatory | Cellular: 25% ↑ WBC | Cellular: 40% ↑ Neutrophil, 50% ↓ | Cellular: 50% ↑ Neutrophil, 50% ↓ | Cellular: 230% ↑ Neutrophil, 60% ↓ |
|  | Cytokine: = ~3x IL-6, IL1ra | Lymphocytes, Monocytes: biphasic | Lymphocytes, Monocytes: biphasic | Lymphocytes, Monocytes: biphasic |
|  | Transcriptomics: Not known | ↓ 60% (3hrs) ↑ 50% (6hrs) | ↓ 30% (3hrs) ↑ 35% (6hrs) | ↓ 60% (3hrs) ↑ 50% (6hrs) |
|  |  | Cytokine: +~3x IL-6, IL1ra, 50% ↑ IL-10 | Cytokine: +~6x IL-6, ~1.3x TNF-α, +/- IL-10 | Cytokine: +++ >10x IL-6, IL1ra, TNF, IL-8, ~5x IL-10 |
|  |  | Transcriptomics: well-described | *Transcriptomics: described | Transcriptomics: TLR-4 described |

The pharmacokinetics of IFN-β may display some variations that will depend on the administration route and dosing. Overall, in healthy individuals, IFN-β has a half-life of 4-5 hr and bioavailability of 30-50%, reaching peak concentrations within 1-8 hr post-injection (Hu et al., 2016; Salmon et al., 1996). Our results show that Plasma IFN-β concentrations increased dramatically by ∼15-fold at 4 hr and ∼9-fold at 6 ½ hr post-injection. Participants also reported a significant rise in side effects starting from 3 hr after administration.

Our findings demonstrate that IFN-β administration as an in-vivo experimental medicine model induces a strong interferon-related and inflammasome-related gene signature, resembling the immune response to infection. Differential expression analysis revealed significant upregulation of genes involved in the interferon pathway, including ISG15, IFIT3, IFIH1, IFI35, DDX58, IRF7, IFI16, IFITM1, and IFNGR2. Additionally, cytosolic nucleic acid sensors such as TLR7 and cGAS, known to trigger IFN-β upon activation, were also upregulated (Costa Franco et al., 2018; Saitoh et al., 2017). Alongside this interferon response, genes regulating the inflammasome pathway, including IL1B, IL18R1, NOD1, NOD2, NLRC4, and NEK7, were upregulated, highlighting activation of the NLRP3 inflammasome. The upregulation of CASP1, CASP4, and CASP5 further suggests both canonical (CASP1) and non-canonical (CASP4, CASP5) inflammasome activation. These results provide new insights into how IFN-β administration mimics the host response to viral infection and triggers an immune response, reinforcing its relevance as a model for studying inflammation.

Consistent with previous findings using LPS and typhoid vaccine immune challenges (De Marco et al., 2022, 2023; Eisenberger et al., 2010; Harrison et al., 2015), IFN-β induced similar transient changes in mood, fatigue, and sickness symptoms. These changes were measured using self-rating questionnaires POMS, fVAS and SicknessQ, commonly used to assess the impact of systemic inflammation on mood and fatigue. This provides further support for using IFN-β as a novel acute inflammatory challenge (see Table 5)

**Table 5.**
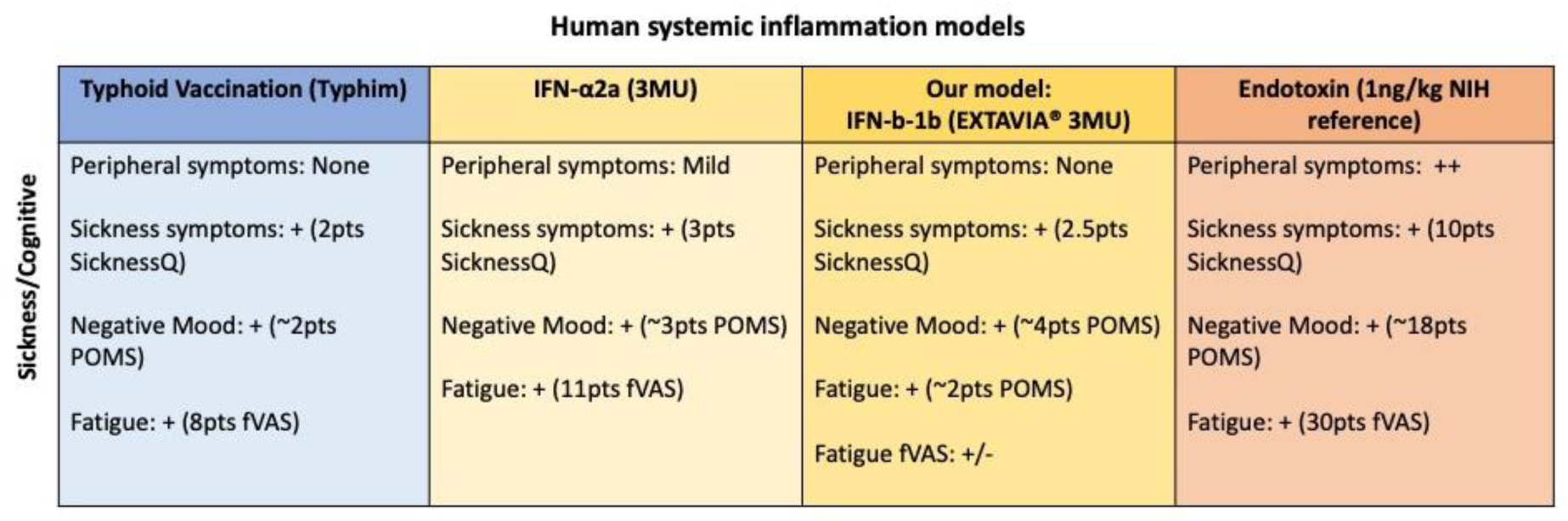
Comparison of the IFN-β model with typhoid, IFN-a, and Endotoxin challenges for the sickness response.

One primary goal of the study was to assess if age influences the effects of IFN-β on physiological, behavioural, and cognitive responses. Given that older individuals often experience more pronounced behavioural responses to infections, and severe infections can lead to lasting cognitive decline (Iwashyna et al., 2010), we hypothesized that inflammation would differentially impact older individuals. Specifically, we expected to observe notable differences in the progression of physiological and immune responses.

No significant condition × age interactions were found for physiological, behavioural or immune markers. Despite this, time × age interactions for IL-6 or condition x age interaction trend on lymphocytes, suggest that IFN-β might still reveal age-related differences. Further studies incorporating larger samples might enhance the detection of subtle interaction effects.

Immune senescence and inflammageing may explain why older individuals often exhibit less efficient immune responses (Franceschi et al., 2018; Pawelec, 2012) Although older age is linked to higher inflammation markers (Cohen et al., 2003; Walston et al., 2002), the exact impact of age on inflammation-induced changes remains unclear. Evidence from sepsis studies indicates a delayed resolution of inflammation in older adults (Bruunsgaard et al., 1999; Kale et al., 2010). In this study, older participants self-reported more intense symptoms post-session, suggesting a potential prolonged inflammatory response. However, practical challenges such as participant burden and logistical constraints limited the feasibility of extended follow-up periods. Further research is needed to elucidate the precise mechanisms underlying age-related differences in inflammation and immune responses. Longitudinal studies with more extensive follow-up periods would warrant deeper insights into the temporal dynamics of inflammation in older adults.

Differences in IFN-β absorption and metabolism between age groups may also account for the observed results. At 4 hr post-challenge, young participants had ∼30% higher circulating IFN-β compared to older individuals, coupled with faster elimination rates. This indicates that older participants may absorb less IFN-β but retain it longer, potentially affecting cytokine responses and the progression of inflammation differently. Such differential metabolism may influence age-related variations in inflammatory responses. Overall, while the study did not find significant age-related differences in IFN-β responses, the observed trends and potential metabolic differences warrant further investigation to fully understand how age affects inflammatory responses.

In conclusion, this study is pioneering in its application of IFN-β within an acute experimental inflammation model. It also represents the first investigation into the effects induced by age-related inflammation. The significant main effects of IFN-β observed in the physiological, behavioural and immune response show that the model can be used as a minimally invasive and effective experimental design able to induce transient changes in systemic inflammation in healthy individuals. As IFN-β elicits a mild but robust response (more effective than the typhoid model but not as strong and invasive as endotoxin challenges), It is suitable for wider experimental applications and can be used in more challenging and complex populations.

## Notes

### Competing Interest Statement

The authors have declared no competing interest.

